# Substance P and adenosine signaling pathways regulate exosomal sorting of miR-21 in colonic epithelial cells

**DOI:** 10.1101/2022.03.30.486482

**Authors:** Sameena Wani, Ivy Ka Man Law, Amy K. Bugwadia, Jill M. Hoffman, Charalabos Pothoulakis

## Abstract

**Background & Aims:** MicroRNAs (miRNAs) are transported in body fluids within exosomes, and exosomal-miRNA sorting is highly selective. In colonic epithelial cells (CECs), Substance P-neurokinin-1-receptor (SP/NK1R) signaling regulates miR-21 sorting into secreted exosomes. Here, we studied the molecular mechanisms driving miR-21 sorting into colonic epithelial exosomes (CEEs) under SP-stimulation.

**Methods:** We performed studies with human colonic epithelial NCM460 cells overexpressing neurokinin-1-receptor (NCM460-NK1R) and in intestinal-epithelial-specific NK1R knockout mice. SP-regulated gene targets were validated by real-time polymerase chain reaction (RT-PCR) and immunoblotting. Small non-coding RNAs (sncRNAs) were isolated from NCM460-NK1R cells and secreted exosomes, and 3′-end adenylated and 3′-end uridylated fractions were separated. Cellular and exosomal sncRNA fractions were processed for miR-21 and miR-1307-3p expression and 3′-end adenylation to uridylation (A/U) ratios using RT-PCR. Pharmacological inhibition studies in NCM460-NK1R cells were performed in the presence of the Adenosine A2B receptor (ADORA2B) antagonist, PSB-1115, followed by RT-PCR and immunoblotting. Mass spectrometry validated in silico interacting partners of miR-21 in CECs.

**Results:** In CECs, miR-21 is predominantly polyuridylated under SP-stimulation and preferentially sorted to CEEs. SP/NK1R signaling activation upregulates extracellular adenosine (ADO) signaling via ADORA2B concomitant with reduced ADO uptake via epithelial-specific equilibrative nucleoside transporter-2 (ENT-2). Mass spectrometry and immunoblotting revealed upregulation of TUT7 in SP-stimulated CEEs. Knockdown of TUT7 decreased both TUT7 and miR-21 content in these exosomes. Pharmacological inhibition of ADORA2B in CECs resulted in decreased TUT7 and miR-21 in CEEs, regardless of SP stimulation.

**Conclusions:** SP/NK1R coupling in CECs activates ADORA2B and its downstream signaling cascade which mediates TUT7/miR-21 interaction and subsequent miR-21 polyuridylation and exosomal export.

**Synopsis:** Substance P-neurokinin-1 receptor signaling regulates terminal uridyl transferase7-mediated predominant 3’-end polyuridylation and exosomal recruitment of miR-21 in human colonic epithelial NCM460 cells overexpressing NK-1R via activation of Adenosine A_2B_ receptor and its downstream signaling cascade. Because Adenosine A_2B_ receptor signaling is involved in IBD pathogenesis, it could therefore be exploited as a pharmacological target for the therapeutic benefits in IBD.

## Introduction

Exosomes are phospholipid-enclosed nanovesicles secreted in response to cell-extrinsic and cell-intrinsic signals under normal and pathological conditions^1,2^. Exosomes encapsulate bioactive molecules from parent cells and exosomal miRNAs represent the most abundant exosomal cargo molecules transferred in active form to adjacent cells or to distant organs^1,3^. MiRNAs are an endogenous group of small non-coding RNAs (sncRNAs) that regulate gene expression post-transcriptionally^4,5^. MiRNAs are sorted to exosomes during multivesicular body (MVB) formation and provide a snapshot of the physiological state of the donor cell^2,6^. Exosome-associated miRNAs mediate crosstalk between cells in both physiological and pathological conditions, including cancer^7–9^ and inflammatory bowel diseases (IBD)^10–14^. MiRNA sorting into exosomes is highly selective and only a subset of cellular miRNAs is encapsulated by exosomes. MiRNAs undergo a series of post-transcriptional modifications, including 5′-end phosphorylation, 3′-end adenylation or uridylation, and terminal nucleotide deletion which possibly contribute to their nanovesicle sorting^15–18^. MiRNA variants with polyadenylated 3′-end sequences are more enriched intracellularly whereas miRNA isoforms with polyuridylated 3′-end sequences are overrepresented in exosomes^16,19,20^. Of note, the RNA-dependent terminal nucleotidyl transferases (TENTs), especially terminal uridyl transferases (TUTases), TUT7 and TUT4, preferentially uridylate a selective subset of miRNAs in cells^15,21–24^. Terminal uridyl transferase7 (TUT7) is not only capable of oligo-uridylating miRNAs but also facilitates miRNA degradation by exosomes, suggesting a pivotal role in exosomal sorting of miRNAs^20,24–26^. A recent study by Wani et al demonstrated that differential distribution of adenosine kinase (ADK) between cells and exosomes modulates the cellular adenosine pool which, in turn, affects the 3′-end adenylation to uridylation ratio of miRNAs and subsequent exosomal recruitment^16^. With the loss of the adenosine pool within cells, adenylated miRNA variants are replaced by uridylated miRNA isomers (i.e., isomiRs). Intriguingly, adenosine levels in different cell types, including colonic epithelial cells (CECs), are regulated by a coordinated network of intestinal epithelium-specific Adenosine A_2B_ receptor (ADORA2B) and the intestinal epithelium-specific equilibrative nucleoside transporter-2 (ENT2)^27–31^. Coupling of adenosine (ADO) to its epithelial-specific receptor ADORA2B maintains epithelial barrier function and mucosal protection during acute colitis^32,33^. Moreover, accumulating evidence suggests that adenosine A_2B_ receptors in the intestinal epithelium are upregulated in IBD and are important in the development of inflammation and IBD^34–37^.

Substance P (SP) is an 11 amino acid neuropeptide/hormone which binds with high affinity to the 407 amino acid G protein-coupled neurokinin-1 receptor (NK-1R) to modulate a myriad of biological functions^11,38,39^. Elevated levels of SP as well as upregulated expression of NK-1R, are reported in the colons of patients with ulcerative colitis (UC) and Crohn’s disease (CD), suggesting a role in IBD pathogenesis^13,39–41^. SP is secreted by many cell types including CECs^13,38,42^, where it couples with NK-1R and elicits proinflammatory signaling^13,38,43,44^ as well as promotes proliferation^13,45^ through activation of MAPK, AKT, and NF-κB signaling cascades and colonic inflammation^13,39^. Moreover, SP activates proinflammatory signaling pathways via NK-1R signaling, including nuclear factor (NF-κB)^43,44^ and c-Jun N-terminal kinase^44,46^ in different cellular systems, including CECs^38,44,47^. Our previous studies have shown that the SP-NK-1R complex regulates differential expression of miRNAs in CECs^13,44^, underscoring a possible epigenetic role within the intestinal epithelium. SP stimulation significantly upregulates several microRNAs, including miR-21 in CECs^13,44^, while it promotes both exosome biogenesis and exosomal shuttling of miR-21 from CECs^13^. Further, internalization of miR-21-enriched exosomes by the target/neighboring IECs implicate their functional significance. However, the molecular mechanism responsible for selective miR-21-exosomal recruitment in CECs remains unknown. As both ADORA2B and SP-NK1R are significantly upregulated in colonic inflammation and IBD, we investigated if these signaling pathways could be related. In the present study, we provide evidence that crosstalk between SP-NK-1R and ADO-ADORA2B signaling pathways orchestrates exosomal sorting of miR-21 in CECs. We demonstrate that SP stimulation of CECs regulates molecular pathways associated with post-transcriptional 3’-end modification of miRNAs contributing to their localization into colonic epithelial exosomes (CEEs). We also identify a role for proinflammatory SP/NK-1R signaling in ADORA2B activation and subsequent TUT7-mediated miR-21 polyuridylation and exosomal export in human CECs.

## Results

### SP-NK1R signaling regulates miR-21 exosomal export by downregulating miR-21-3’-end-polyadenylation

Previously we have reported that SP-NK1R signaling activation regulates miRNA expression profile in human colonic epithelial cells overexpressing NK-1R (NCM460-NK1R) in addition to eliciting cell proliferation and proinflammatory responses in these cells^13,38,40,43^. Exosome production and intracellular levels of miR-21 and miR-1307-3p were upregulated during SP stimulation in human NCM460-NK1R cells^13,44^. We also showed that expression of CD9 was significantly increased in NCM460-NK1R exosomes. In the present study, we further characterized the exosomes produced by NCM460-NK1R cells by examining CD9 and A33 expressions in NCM460-NK1R exosomes under SP stimulation (100nM, 6hr). In contrast to CD9, which is a ubiquitous exosome marker^50^, A33 is an intestinal epithelial-specific exosome marker^55^. Here we found that expression of both CD9 and A33 proteins were significantly increased in NCM460-NK1R exosomes under SP stimulation (100nM, 6hr) compared to vehicle controls (0.1% TFA). However, CD9 expression was more significant in SP-stimulated exosomes as compared to the intestinal epithelial-specific exosome marker, A33. As depicted in Fig.1A, our immunoblot analysis demonstrates increased expression of CD9 (3.533±0.08, *p*<0.0001, Fig.1A) in SP-stimulated NCM460-NK1R exosomes; while expression of A33 was increased in SP-stimulated cells (1.34±0.09, *p*=0.0002, Fig.1A) compared to control (0.1% TFA). We also quantified miR-21 and miR-1307-3p expression levels in these cells, in the presence or absence of SP stimulation (100nM, 6hr). Similar to our previous findings^44^, we observe a striking increase in the expression levels of intracellular miR-21 (5.403±0.05, *p*<0.0001, Fig.1B) and miR-1307-3p (3.89±0.1, *p*<0.0001, Fig.1B) compared to controls upon SP-NK1R signaling activation in NCM460-NK1R cells.

**Figure 1.**
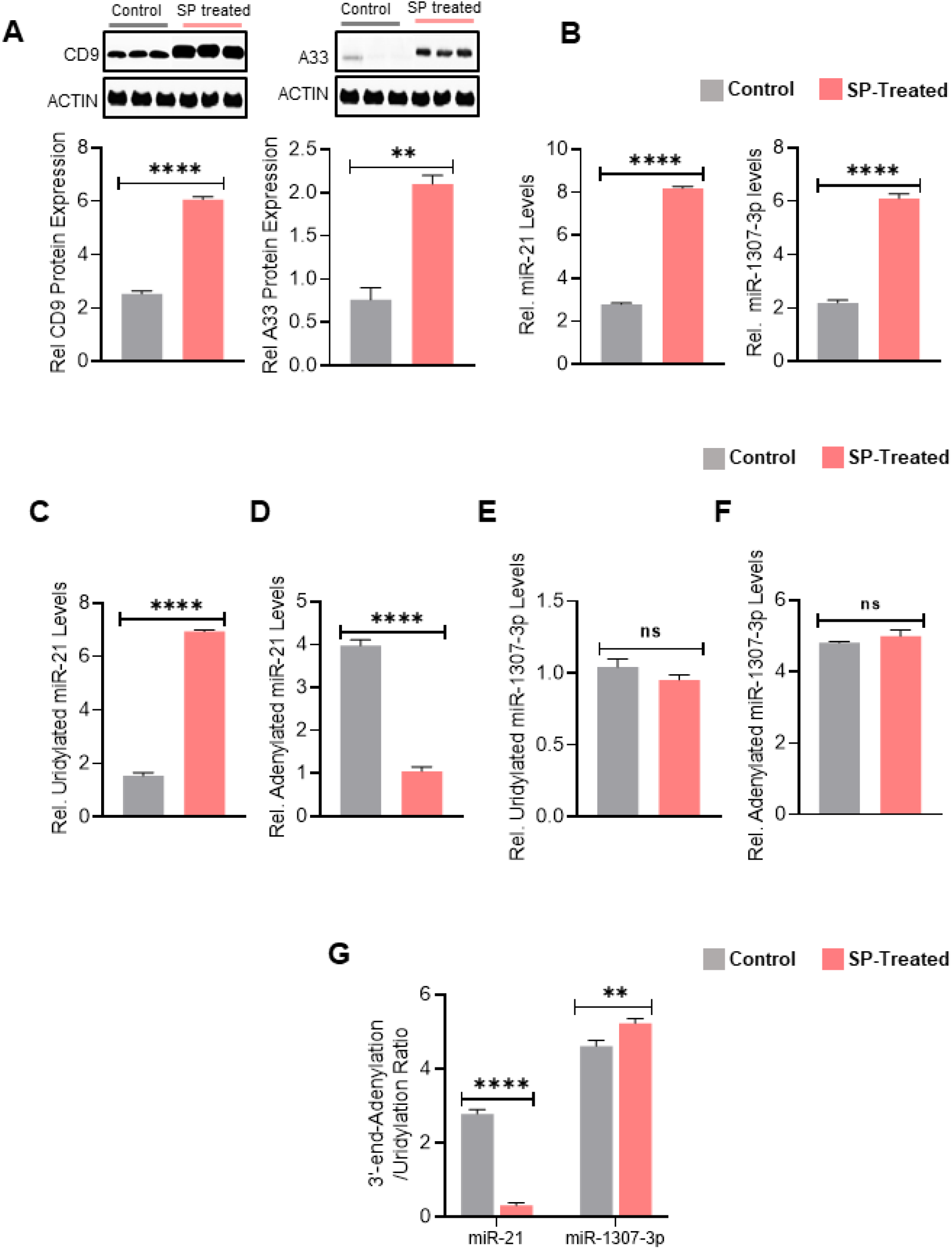
SP-NK1R signaling activation regulates exosome production and miR-21 3’-end uridylation and exosomal sorting in colonic epithelial cells. **(A)** Exosomes released by SP-treated NCM460-NK1R cells (100nM, 6hrs) and their respective controls (0.1%TFA) were collected and analyzed for the expression of exosome biomarkers, including CD9 and A33 by immunoblotting. Actin was used as a loading control **(B)** Cellular expression of miRNAs, miR-21 and miR-1307-3p, under SP-stimulation (100nM, 6hrs) vs Vehicle control (0.1%TFA). Levels of polyuridylation (in SP-stimulated exosomes vs vehicle control) of **(C)** miR-21 and **(E)** miR-1307-3p. Levels of polyadenylation (in SP-treated NCM460-NK1R cells vs vehicle control) of **(D)** miR-21 **(F)** miR-1307-3p **(G)** Results are represented as 3’-end adenylation/uridylation ratio for the miRNAs, miR-21 and miR-1307-3p in NCM460-NK1R cells (with or without SP-stimulation). Ratio was analyzed to ascertain the predominant post-transcriptional modification of miRNAs in these cells under SP-NK1R signaling activation. Gene expression analysis was done using RT-PCR and expression was calculated relative to GAPDH or U6. Experiments were performed three independent times and all experiments were carried out in triplicate. All bar graphs represent mean and error bars are SD (mean ± SD), n.s. non-significant, \**p*<0.05, \*\**p*<0.01, \*\*\**p*<0.001, \*\*\*\**p*<0.0001 by unpaired t-test.

MiR-21 is predominantly exported to exosomes while miR-1307-3p remains intracellular upon SP stimulation^13^. To examine our hypothesis that miR-21 export to intestinal epithelial exosomes is dependent on levels of 3’-end adenylation and uridylation, we compared levels of 3’-end adenylation and uridylation of intracellular miR-21 and miR-1307-3p, in the presence and absence of exogenous SP-NK1R signaling activation in human colonic epithelial NCM460-NK1R cells. Intracellular miRNAs with 3’-end adenylation and uridylation were collected as separate fractions from NCM460-NK1R cells treated with SP (100nM, 6hr) and control (0.1% TFA) and then the levels of miR-21 and miR-1307-3p were analyzed in each polyadenylated and polyuridylated miRNA fractions. As shown in Fig.1C, SP stimulation preferentially increased 3’-end uridylation of miR-21 (5.390±0.07, *p*<0.0001, Fig.1C) while decreasing miR-21-3’-end adenylation (2.925±0.06, *p*<0.0001, Fig.1D). In contrast, the levels of 3’-end-uridylation (0.090±0.03, *p*=0.085, Fig.1E) and -adenylation of miR-1307-3p (0.183±0.10, *p*=0.15, Fig.1F) did not change as a result of SP treatment. Overall, SP stimulation led to a significant reduction in the adenylation to uridylation (A/U) ratio of miRNA, miR-21 (2.473±0.07, *p*<0.0001, Fig.1G), while this ratio was increased for the SP-upregulated miRNA, miR-1307-3p (0.613±0.10, *p*=0.0054, Fig.1G). Taken together, our results suggest that SP-NK-1R-associated export of miR-21 to exosomes is mediated by increased uridylation and decreased adenylation at the 3’-end sequence of miR-21 in human CECs.

### SP-NK1R signaling activation regulates adenosine uptake in colonic epithelial cells in vitro and in vivo

To understand the molecular process by which SP-NK1R signaling affects miR-21 post-transcriptional modification, genes associated with adenosine uptake were examined in NCM460-NK1R cells. Amongst the equilibrative nucleoside transporters ENT1 and ENT2, ENT2 is more selectively expressed in colon than ENT1^27^. Our results showed that SP stimulation (100nM, 6hr) in intestinal epithelial cells (IECs) significantly reduced ENT2 expression (2.173±0.10, *p*<0.0001, Fig.2A) while ENT1 expression remained unchanged (0.066±0.05, *p*=0.29, Fig.2A) regardless of SP-NK1R signaling activation. These findings are in line with previous studies^27,56,57^, and suggest that ENT2 and not ENT1 is a major transporter for adenosine in IECs. We also examined IL8 expression, a hallmark of SP stimulation in NCM460-NK1R cells^38,43^. We showed that concurrent with a decrease in ENT2 expression (2.173±0.10, *p*<0.0001, Fig.2A), a significant increase in IL8 expression was also observed (6.913±0.10, *p*<0.0001, Fig.2A), suggesting activation of SP-NK1R signaling in these cells.

**Figure 2.**
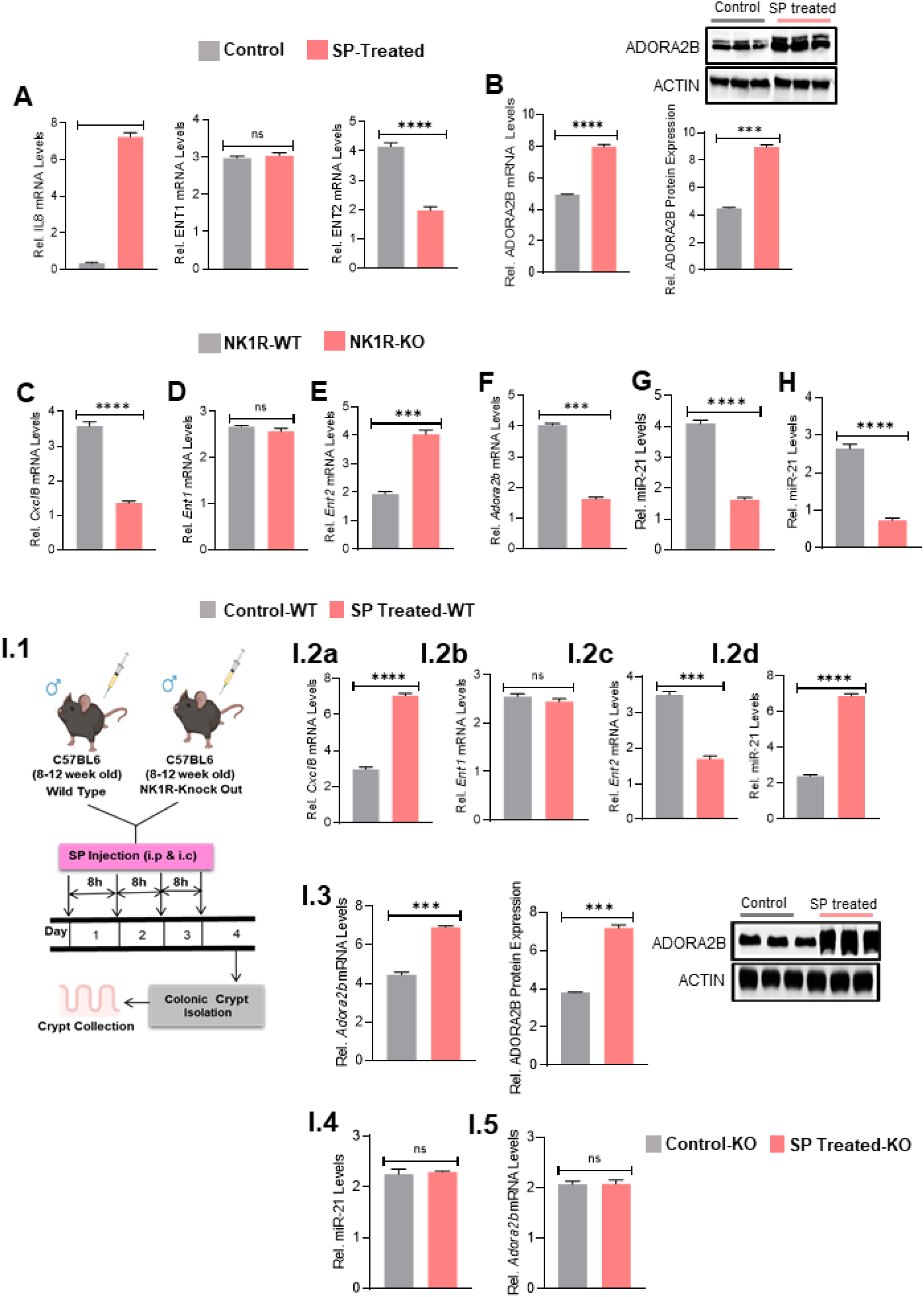
SP stimulation affects adenosine transporter and receptor expressions in colonic epithelial cells. **(A)** Gene expression of IL8, ENT-1 & −2 in SP-treated NCM460-NK1R cells vs vehicle controls (0.1%TFA) were examined using RT-PCR. **(B)** Gene and protein expressions of ADORA2B in NCM460-NK1R cells treated with SP and their control counterparts (0.1%TFA) was analyzed using RT-PCR and immunoblot, respectively. Gene expression of **(C)***Cxcl8*, **(D)** *Ent-1*, **(E)** *Ent-2* and **(F)** *Adora2b* in colonic tissues collected from Vil1-cre;NK1R^+/+^ and Vil1-cre;NK1R^fl/fl^ mice (*n*=8 per group). Expression of miR-21 **(G)** in colonic epithelial units/ crypts (*n*=8 per group) and **(H)** colonic tissues (*n*=8 per group) derived from Vil1-cre;NK1R^+/+^ mice and Vil1-cre;NK1R^fl/fl^. Gene expressions of *Cxcl8* **(I.2a)**, *Ent1* **(I.2b)**, *Ent2* **(I.2c)** and miR-21 **(I.2d)** in colonoids derived from Vil1-cre;NK1R^+/+^ mice treated with SP and its vehicle control (0.1%TFA). **(I.3)** Gene and protein expressions of ADORA2B in colonoids derived from Vil1-cre;NK1R^+/+^ mice treated with SP and its vehicle control (0.1%TFA). Gene expressions of miR-21 **(I.4)** and *Adora2b* **(I.5)** in colonoids derived from Vil1-cre;NK1R^fl/fl^ mice treated with SP and its vehicle control (0.1%TFA). Results represent *n* =8 mice per group from 2 independent experiments. Actin was used as a loading control for western blot analysis. Gene expression analysis was done using RT-PCR and expression was calculated relative to GAPDH or U6. Experiments were performed three independent times and all experiments were carried out in triplicate. All bar graphs represent mean and error bars are SD (mean ± SD), n.s. non-significant, *p<0.05, **p<0.01, ***p<0.001, ****p<0.0001 by unpaired t-test.

Studies have shown that ADORA2B, the predominant receptor for adenosine expressed on IECs^27,28,30,37^, is upregulated in IBD^27,34^ and its signaling elicits a protective response in the intestinal mucosa^30^ and promotes intestinal epithelial barrier repair during colitis^27,30,34^. In our study, we examined the expression of intestinal epithelial-specific adenosine A_2B_ receptors in SP stimulated (100nM, 6hr) NCM460-NK1R cells and murine colonic epithelial cells isolated from C57BL6 wild type mice (n=8 for each group) following treatment with SP (72μg/mouse, 100μL). As shown in Fig.2B, our results clearly indicate a significant increase in ADORA2B gene (3.045±0.08, *p*<0.0001, Fig.2B) and protein expression (4.469±0.09, *p*=0.0004, Fig.2B), in human colonic epithelial NCM460-NK1R cells. Taken together, these data suggest that SP-NK1R signaling activation in human colonic epithelial NCM460-NK1R cells results in a significant reduction of adenosine uptake via its transporters (in particular, ENT2 in CECs) resulting in decreased intracellular adenosine levels and increased extracellular adenosine signaling through ADORA2B.

To further examine the association between SP-NK1R signaling and expression of *Ent2* and *Adora2b* in colonic epithelium, colonic tissues from intestinal epithelial-specific NK-1R knockout mice (Vil1-cre;NK1R^fl/fl^) and controls (Vil1-cre;NK1R^+/+^) were collected and gene expression of *Ent2* and *Adora2b* were examined. As shown in Fig.2C, *Cxcl8* expression, the mouse homolog to IL8, was significantly reduced in Vil-cre;NK1R^fl/fl^ mice when compared to wild type (Vil1-cre;NK1R^+/+^)(2.223±0.07, *p*<0.0001, Fig.2C). In line with our *in vitro* results, NK-1R deficiency significantly increased *Ent2* expression in NK-1R-KO mice (2.087±0.10, *p*=0.0004, Fig.2E) while *Ent1* expression remained unchanged (0.100±0.03, *p*=0.06, Fig.2D). Intriguingly, we also observed a significant reduction of *Adora2b* expression (2.403±0.04, *p*=0.0001, Fig.2F) in the colonic epithelia of NK-1R-KO mice. Importantly, miR-21 levels were dramatically repressed in colonic epithelial units/crypts (2.477±0.07, *p*<0.0001, Fig.2G) as well as colonic tissues (1.910±0.07, *p*<0.0001, Fig.2H) taken from Vil-cre;NK1R^fl/fl^ mice (n=8), suggesting that SP-NK1R signaling regulates expression of *Ent2, Adora2b* and miR-21 in the colonic epithelium.

Lastly, colonic epithelial stem cells were isolated from Vil1-cre;NK1R^fl/fl^ and their wild type littermates (n=8 for each group) and cultured into colonoids (*ex vivo*). Colonoids were treated with SP (100nM, 6hr); and SP-NK1R signaling activation was confirmed by significantly increased *Cxcl8* expression, the mouse homolog to IL8 (4.077±0.10, *p*=<0.0001, Fig.2-I.2a). Results from gene expression analysis showed that SP stimulation not only caused a significant reduction of *Ent2* gene expression (1.810±0.07, *p*=0.0005, Fig.2-I.2c) in these colonoids but also a simultaneous significant increase of *Adora2b* at both the transcriptional (2.483±0.08, *p*=0.0004, Fig.2-I.3) and translational levels (3.380±0.09, *p*=0.0002, Fig.2-I.3). Again, no change in *Ent1* gene expression was observed in the Vil1-cre;NK1R+/+-derived colonoids regardless of SP treatment (0.100±0.04, *p*=0.10, Fig.2-I.2b). Interestingly, expression levels of miR-21 in colonoids derived from Vil1-cre;NK1R^+/+^ mice treated with SP were significantly increased (4.450±0.09, *p*<0.0001, Fig.2-I.2d) compared to colonoids treated with vehicle controls (n=8 for each group). Additionally, gene expression analysis of miR-21 (0.043±0.05, *p*=0.49, Fig.2-I.4) and *Adora2b* (0.006±0.05, *p*=0.91, Fig.2-I.5) in Vil1-cre;NK1R^fl/fl^ -derived colonoids treated with or without SP, showed no significant effect on miR-21 or *Adora2b* expression which further supports the concept that SP-NK1R regulates miR-21 and *Adora2b* signaling in colonic epithelial cells. Overall, our results provide further evidence that SP-NK1R signaling activation potentially reduces adenosine uptake via its transporters (ENT1 & ENT2) while simultaneously increasing extracellular adenosine signaling via ADORA2B in colonic epithelial cells *in vivo*.

### SP-NK1R signaling activation promotes TUT7/CREB interaction with miR-21 promoter region mediating miR-21 polyuridylation and exosomal sorting

cAMP response element-binding protein (CREB) is not only a proto-oncogenic transcription factor but also a transcriptional activator known to enhance the expression of target genes^58^. Previous studies have shown that miR-21 promoter harbors four conserved CREB-binding sites^59^ and that CREB regulates miR-21 promoter activity^59^. To provide mechanistic insights into SP-NK1R signaling, ADO-ADORA2B signaling and miR-21 expression as well as its exosomal trafficking, bioinformatics analysis was performed to identify potential genomic regulators associated with miR-21. Results from open-access web-based bioinformatics tools, including TransmiRv2.0^60^ and RNA-Protein Interaction Prediction (RPISeq)^51^, suggested that cAMP responsive element-binding protein (CREB) and Terminal uridylyl transferase 7 (TUT7) are two candidates for miR-21 promoter binding (Fig.3A.a, c). Our TransmiRv2.0 analysis revealed that CREB binds miR-21 from region 59837682 to 59837697 on chromosome 17 (>H.sapiens; range=chr17:59837682-59837697 5’pad=0 3’pad=1; strand=(-) GGGAGGCACCTCCCA) (Fig.3A.a). Also, our RPISeq analysis revealed a very strong interaction between miR-21 and TUT7 with an interaction probability of 0.85^####^ (Prediction using SVM classifier) (Fig.3A.c). Additionally, using online software tools including, TFSEARCH^54^, TRANSFAC^53^ and JASPAR^52^ for Transcription factor binding sites, we found a putative binding site for CREB in the promoter region of TUT7 gene (Fig.3A.b).

**Figure 3.**
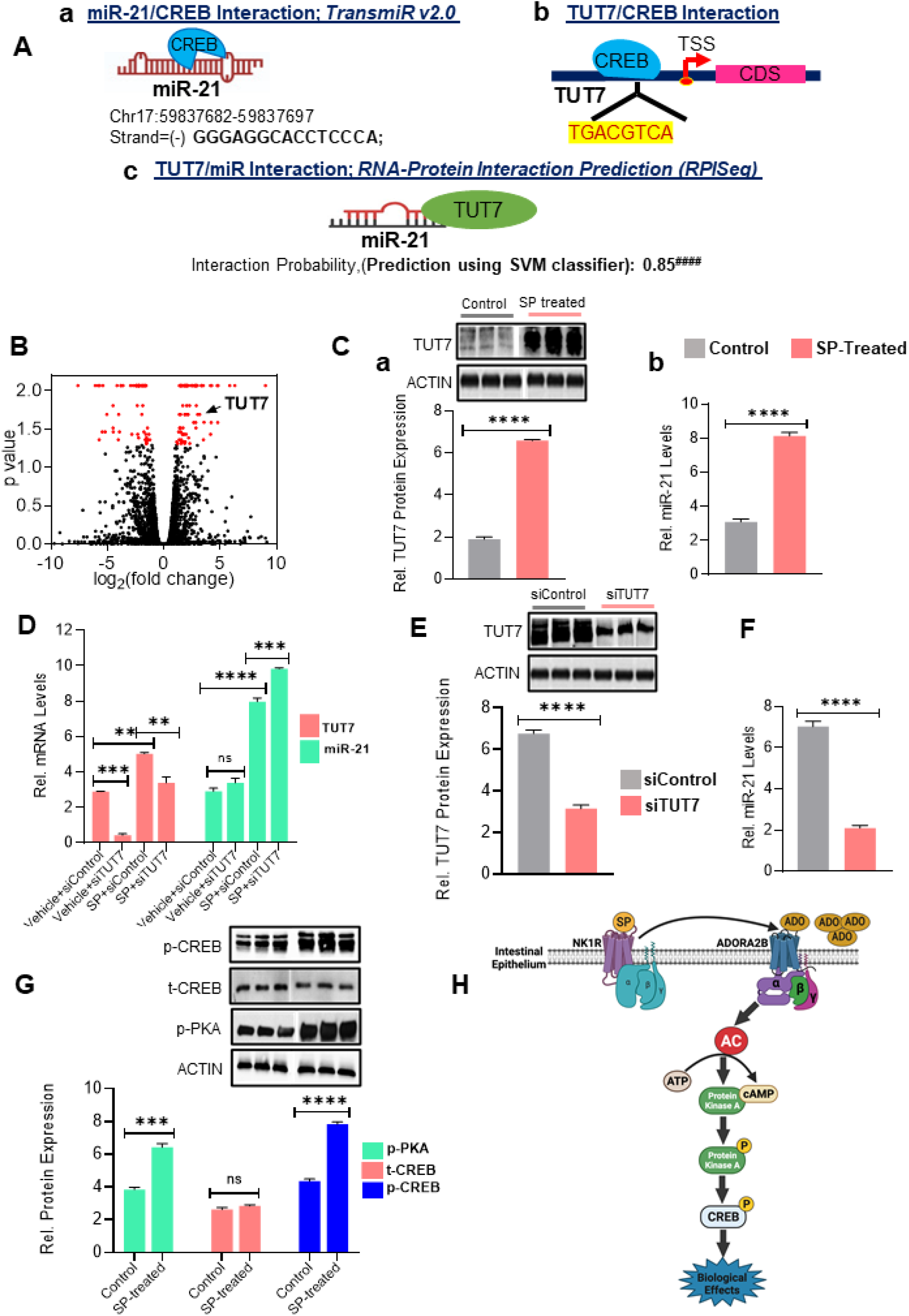
TUT7 binds miR-21 and regulates miR-21 3’-end post-transcriptional modification and its exosomal sorting in human colonic epithelial cells. **(A)** Bioinformatics analysis on the potential interaction of CREB and TUT7 on miR-21 promoter sequence. **(B)** Volcano plot depicting differential protein expression in exosomes secreted by SP-treated NCM460-NK1R cells (100nM,6hrs) and their respective vehicle controls (0.1%TFA) analyzed by mass spectrometry. The volcano plot **(B)** represents the most highly differentially expressed exosomal proteins, red dots represent the up-regulated proteins and green dots represent the down-regulated ones. **(C.a)** Representative immunoblot confirming mass spectrometry results. Expression of TUT7 **(C.a)** and miR-21 **(C.b)** in the same samples was confirmed using immunoblot and RT-PCR, respectively. **(D)** Gene expression of TUT7 and miR-21 was examined in NCM460-NK1R cells transfected with siTUT7 and its control, with or without SP stimulation. **(E)** Protein expression of TUT7 and **(F)** miR-21 expression were analyzed in exosomes released by NCM460-NK1R cells transfected with siTUT7 and its control. **(G)** Western blot analysis of total and phosphorylated CREB as well as phosphorylated PKA in NCM460-NK1R cells treated with SP (100nM,6hrs) or the vehicle control (0.1%TFA). Actin was used as a loading control for western blot analysis. Gene expression analysis was done using RT-PCR and expression was calculated relative to GAPDH or U6. Experiments were performed three independent times and all experiments were carried out in triplicate. All bar graphs represent mean and error bars are SD (mean ± SD), n.s. non-significant, *p<0.05, **p<0.01, ***p<0.001, ****p<0.0001 by unpaired t-test and One-way ANOVA (Tukey’s post hoc test).

Interestingly, mass spectrometry analysis on exosomal proteins isolated from SP-stimulated human colonic epithelial NCM460-NK1R cells (100nM,6hr) and those from treatment controls (0.1%TFA, vehicle control) further validated our in-silico results. Fig.3B is the representative volcano plot depicting differential exosomal protein expression from NCM460-NK1R cells with or without SP stimulation as analyzed by mass spectrometry. These results revealed that an increased amount of TUT7 protein is exported to exosomes upon SP stimulation in NCM460-NK1R cells, when compared to TUT7 levels in exosomes collected from cells under control treatment (0.1%TFA). These results were further validated by immunoblotting as indicated in Fig.3C.a (4.697±0.06, *p*<0.0001, Fig.3C.a). Of note, similar to our previous studies^13^, results from the current study also revealed that miR-21 levels were increased significantly in these exosomes (5.033±0.14, *p*<0.0001, Fig.3C.b), besides, TUT7, suggesting that TUT7 plays a pivotal role in the exosomal transport of miR-21 in human colonic epithelial cells. We next examined cellular and exosomal levels of miR-21 in NCM460-NK1R cells deficient in TUT7, in the presence and absence of SP stimulation (100nM,6hr). Transfection with si-TUT7 significantly reduced TUT7 expression in cells treated with SP (100nM,6hr) (1.643±0.15, *p*=0.001, Fig.3D) as well as the control (2.457±0.05, *p*=0.0001, Fig.3D). Interestingly, we noticed that the siTUT7-mediated reduction of TUT7 with concurrent SP-NK1R activation further increased the cellular miR-21 levels in NCM460-NK1R cells compared to siControl treatment (1.863±0.10, *p*=0.0001, Fig.3D). However, no change in miR-21 levels was observed in TFA-treated control cells deficient in TUT7 (0.4767±0.15, p=0.08, Fig.3D). In contrast, TUT7 deficiency caused a significant reduction of both TUT7 (3.623±0.13, p<0.0001, Fig.3E) and miR-21 expression levels (4.937±0.16, p<0.0001, Fig.3F) in exosomes secreted by TUT7-deficient NCM460-NK1R cells compared to control cells (siControl treated).

In addition to our bioinformatic analysis of RNA-protein interaction, previous studies also suggest that CREB is downstream to ADORA2B signaling in many cell types, especially CECs^28,37^. Keeping this in mind, we next examined whether protein kinase A (PKA) and CREB were activated upon SP-NK1R coupling. Our immunoblot analysis (Fig.3G) of lysates collected from SP-treated NCM460-NK1R cells (100nM,6hr) and their vehicle controls (0.1%TFA) showed significantly increased phosphorylation of both PKA (2.573±0.15, *p*=0.0005, Fig.3G) and CREB (3.477±0.12, *p*<0.0001, Fig.3G) proteins. However, expression levels of total CREB (t-CREB) remained unchanged in SP-treated NCM460-NK1R cells vs control cells (0.213±0.08, *p*=0.06, Fig.3G). These results further validate our *in silico* findings showing possible interactions between CREB, TUT7 and miR-21. Overall, our results indicate that SP stimulation in NCM460-NK1R cells not only regulates TUT7 and miR-21 expression but also is pivotal in the regulation of PKA and CREB. SP-induced PKA/CREB expression in NCM460-NK1R cells in turn regulate the expressions of both miR-21 and TUT7. Taken together, these results demonstrate that SP-NK1R signaling activation in human colonic epithelial NCM460-NK1R cells promote miR-21 exosomal recruitment possibly through PKA/CREB activation and subsequent TUT7 interaction with miR-21. TUT7 interaction with miR-21 mediates polyuridylation and exosomal export of miR-21 in NCM460-NK1R cells.

### SP-NK1R signaling activation promotes TUT7-CREB-dependent miR-21 recruitment to exosomes through transactivation of ADORA2B

SP-NK1R signaling leads to transactivation of insulin-like growth factor receptor (IGFR) thereby contributing to both proinflammatory and proliferative responses in colonic epithelium^61^. Based on our results described above, we next examined whether SP-induced PKA/CREB activation and TUT7 and miR-21 exosomal export in colonic epithelial cells was dependent on ADRORA2B receptor activity in these cells. PSB-1115 (100nM,1hr), an ADORA2B receptor antagonist^27^ was used to inhibit ADORA2B signaling (Fig.4A) in human colonic epithelial cells in the presence or absence of SP stimulation (100nM,6hr). Immunoblot analysis showed that ADORA2B inhibition (PSB-1115;100nM,1hr) in NCM460-NK1R cells significantly reduced p-PKA levels in control cells (1.377±0.19, *p*=0.0009, Fig.4B) and SP-treated cells (4.120±0.15, *p*=0.0001, Fig.4B). Similarly, CREB activation (levels of p-CREB) was also downregulated in both vehicle control cells (1.117±0.05, *p*=0.0001, Fig.4B) as well as SP-treated cells (5.683±0.008, *p*<0.0001, Fig.4B) upon ADORA2B inhibition in these cells.

**Figure 4.**
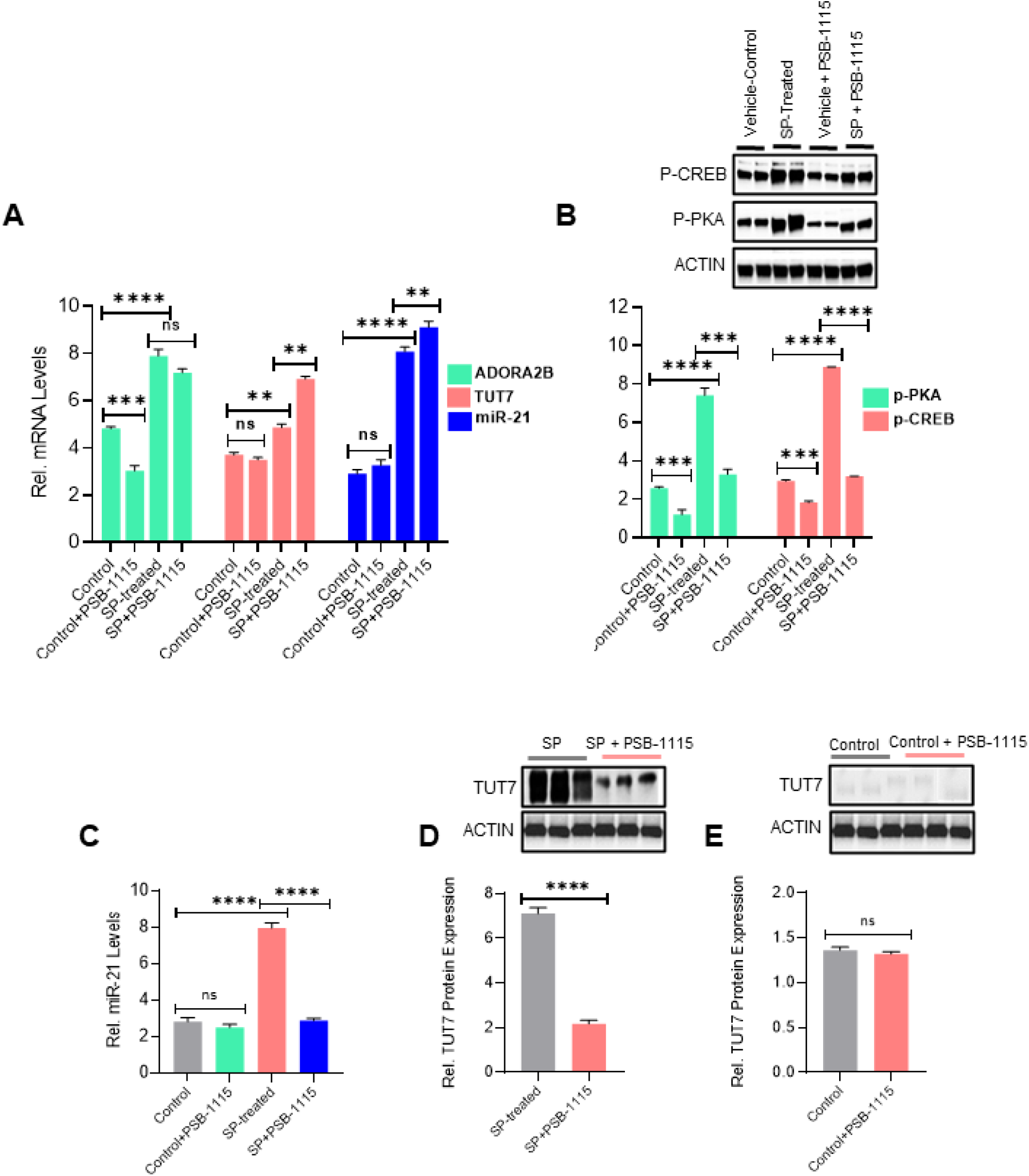
SP-induced ADO/ADORA2B signaling activation triggers intracellular cascade resulting in downstream miR-21-TUT7 binding and miR-21 polyuridylation and exosomal export. **(A)** Gene expression of ADORA2B, TUT7 and miR-21 in NCM460-NK1R cells treated with PSB-1115 (100nM,1hr) or its vehicle control (sterile water) in the presence or absence of SP stimulation. **(B)** Immunoblot analysis of phosphorylated PKA and CREB levels in NCM460-NK1R cells treated with PSB-1115 (100nM,1hr) or its vehicle control in the absence or presence of SP stimulation. **(C)** miR-21 levels in the exosomes secreted by NCM460-NK1R cells under PSB-1115 treatment (100nM,1hr) or its vehicle control in the presence or absence of SP stimulation. **(D&E)** Expression of TUT7 measured by western blot in the exosomes secreted by NCM460-NK1R cells treated with PSB-1115 (100nM,1hr) or its vehicle control in the presence of SP stimulation**(D)** and 0.1% TFA**(E)**. For western blot analysis actin was regarded as the internal reference. Gene expression analysis was done using RT-PCR and expression was calculated relative to GAPDH or U6. Experiments were performed three independent times and all experiments were carried out in triplicate. All bar graphs represent mean and error bars are SD (mean ± SD), n.s. non-significant, *p<0.05, **p<0.01, ***p<0.001, ****p<0.0001 by unpaired t-test and One-way ANOVA (Tukey’s post hoc test).

Subsequent gene expression analysis (Fig.4A) showed that while ADORA2B inhibition did not have significant effects on ADORA2B gene expression after SP stimulation in NCM460-NK1R cells (0.713±0.17, *p*=0.05, Fig.4A), both cellular TUT7 (2.053±0.09, *p*=0.002, Fig.4A) and miR-21 (1.023±0.14, *p*=0.005, Fig.4A) expression levels were significantly increased after ADORA2B inhibition in SP-treated NCM460-NK1R cells. In contrast, ADORA2B inhibition significantly reduced TUT7 (4.933±0.16, *p*=0.0001, Fig.4D) and miR-21 (5.067±0.14, *p*<0.0001, Fig.4C) expression levels in exosomes secreted from SP-treated NCM460-NK1R cells, suggesting that ADORA2B inhibition in NCM460-NK1R cells significantly decreased the export of both TUT7 and miR-21 to the secreted exosomes. Of note, no such changes were observed in the expression levels of either miR-21 (0.320±0.15, *p*=0.12, Fig.4C) or TUT7 (0.043±0.02, *p*=0.17, Fig.4E) in exosomes secreted from SP-treated NCM460-NK1R cells under ADORA2B inhibition. Collectively, our results demonstrate that SP-NK1R signaling in human epithelial NCM460-NK1R cells promote miR-21 export to exosomes by two potential mechanisms, 1) transactivating ADO/ADORA2B signaling, resulting in PKA/CREB/TUT7/miR-21 signaling cascade activation and subsequently, promoting downstream interaction of CREB and TUT7 with miR-21. 2) increasing 3-end uridylation of miR-21 by TUT7 resulting in preferential exosomal sorting of miR-21 in these cells^20,22,25^.

## Discussion

SP, and its high-affinity receptor, NK1R, are key players in the pathophysiology of IBD, a debilitating and chronic inflammatory disorder of the gastrointestinal tract^39,44,47^. Levels of SP and NK1R are elevated in the inflamed tissues in IBD^39–41,47^. Dysregulation of miRNAs is widely observed in many pathophysiological processes, importantly cancer and IBD^12,14,62,63^ implying the importance of miRNAs in cellular homeostasis. MiR-21, a multi-faceted RNA, is involved in myriad cellular biological pathways and can regulate numerous cellular processes, including inflammation^8–10,12,59^. Our previous studies have revealed that SP-NK1R signaling stimulates differential miRNA expression in CECs^11,44^ as well as cell proliferation, apoptosis and inflammatory responses^13,38,43–45^. The results from the current study further confirm our previous findings that SP-NK1R signaling activation in NCM460-NK1R cells promotes exosome biogenesis as indicated by significantly increased CD9 and A33 protein expressions in SP-stimulated exosomes. Consistent with results from our previous studies, we also found that SP-NK1R signaling activation significantly increased intracellular miR-21 and miR-1307-3p levels in CECs but only miR-21 in the presence of SP is preferentially transported to exosomes secreted by CECs. These results suggest that SP-stimulation is pivotal in the exosomal sorting of miR-21 in CECs.

MiRNAs encapsulated in exosomes can be efficiently delivered to recipient cells where they exert functional roles^2,11,50^. Notably, miRNAs with oncogenic and inflammatory roles are aberrantly increased in exosomes^11,13^ and specific miRNAs are enriched in secreted exosomes, suggesting that cells exhibit selectivity in sorting miRNA content^11,17,18,50^. Despite increasing attention towards exosome-incorporated miRNAs as potential biomarkers and therapeutic agents, the molecular mechanisms and/or key players which regulate their selective loading remain unclear. Several recent investigations have revealed that post-transcriptional 3’-end modifications, polyadenylation and polyuridylation of miRNAs, are responsible for their intercellular containment or exosomal enrichment^16,18–20^. Interestingly, we observed a predominant post-transcriptional 3’-end uridylation of miR-21 under SP-stimulation in CECs. However, SP-NK1R signaling activation did not yield effects on the post-transcriptional modification of miR-1307-3p 3’end sequence in CECs. These results indicate that the SP-stimulation in CECs preferentially polyuridylates miR-21 and may contribute to its sorting into the exosomes secreted by these cells. Our results strongly suggest a role of SP-NK1R signaling activation in CECs in regulating the post-transcriptional modification of the 3’-end sequence of miR-21 and its subsequent exosomal export.

The post-transcriptional modification of the 3’-end sequence of miRNAs, miRNA-polyadenylation and -polyuridylation, is largely dependent on the intracellular adenosine pool^16,20^. Wani et al previously demonstrated that miR-2909 inclusion or exclusion from exosomes is linked to the differential distribution of adenosine kinase, which affects cellular adenosine levels and hence miR-2909 post-transcriptional modifications in prostate cancer^18^. Adenosine, an immunomodulatory biomolecule, elicits many pathological and physiological cellular and tissue functions, including inflammation, by binding to four specific GPCRs, A1, A2A, A2B, and A3, (gene names, ADORA1, ADORA2A, ADORA2, ADORA3), whereas the intracellular uptake of adenosine is facilitated by ENTs^27,28,30^. Moreover, ADORA2B is predominantly expressed in intestinal epithelium^27,28,37^ and is upregulated in IBD^27,35–37^. Besides, it’s signaling activation in CECs plays a role in epithelial barrier function and mucosal healing in IBD^27,30^. In the present study, we observed that SP-NK1R signaling activation resulted in significantly decreased ENT2 expression in CECs. In contrast, we observed significant upregulation of intestinal epithelium-specific Adenosine A_2B_ receptor under SP-stimulation *in vivo* as well as *in vitro*. These findings are in line with recently reported studies showing interplay between repression of equilibrative nucleoside transporter2 in intestinal epithelium and enhanced extracellular adenosine signaling ^27,30^. The results from the present study further suggest that interaction of SP with its preferred receptor, NK1R, leads to increased ADO signaling via intestinal epithelium-specific ADORA2B and subsequent activation of its downstream signaling cascade in CECs. Overall, these findings demonstrate evidence for biological crosstalk within the colonic epithelium between two major regulatory signaling pathways in IBD, SP-NK1R and ADO-ADORA2B.

Numerous studies have shown the extensive interactions between RBPs and cellular as well as exosomal shuttle RNAs and the key role of these RBPs in packaging of RNAs (especially miRNAs) into exosomes^11,17^. Terminal uridyl transferase7 (TUT7) (also known as ZCCHC6/TENT3B), an RNA-dependent terminal nucleotidyl transferase (TENTs), mediates the addition of non-templated nucleotides to 3′-end sequence of miRNAs, besides, regulating miRNA biogenesis and maturation^20,22,24,25^. In this study, using online bioinformatic tools, including TRANSFAC and RPIseq databases, we identified a strong interaction between TUT7 protein and miR-21. Mass spectrometry and immunoblot analysis provide strong evidence that SP-NK1R signaling activation results in significant upregulation of TUT7, in colonic epithelial exosomes (CEEs). Hence our results show that SP-NK1R signaling activation upregulates both miR-21 as well as TUT7 in CEEs. Additionally, genetic studies involving the knockdown of TUT7 in CECs resulted in a significant decrease of both TUT7 and miR-21 expression levels in exosomes secreted by these cells, further confirming the TUT7/miR-21 binding and TUT7-dependent export of miR-21 into secreted exosomes. Taken together, these findings indicate that SP stimulation in CECs regulates the potential interaction and co-localization of TUT7 and miR-21 in exosomes released by CECs. Our results further indicate that SP-NK1R signaling activation regulates the expression of TUT7 in the colonic epithelium which has a pivotal role in the post-transcriptional 3’-end modification of miR-21 and its transport between colonic epithelial cells and their secreted exosomes.

Interestingly, pharmacological inhibition of adenosine A_2B_ receptor concomitant with SP stimulation in CECs resulted in a significant decrease in expressions of cellular p-CREB as well as p-PKA and a concurrent decrease in the levels of TUT7 protein and miR-21 in secreted exosomes, regardless of SP-NK1R signaling activation. Additional *in silico* analysis (using TRANFAC & JASPAR databases) demonstrated that CREB is a strong candidate for both TUT7 and miR-21 promoter binding and transactivation. These findings are consistent with previous studies demonstrating that CREB is downstream to ADORA2B signaling (adenosine/ADORA2B/cAMP/PKA/CREB pathway), and the ADO/ADORA2B coupling in CECs results in activation of downstream PKA-CREB signaling pathways^28,37^. Further, the present study demonstrates that NK1R stimulation by SP in CECs leads to increased ARORA2B activation and its downstream signaling cascade, and activated ADORA2B signaling promotes TUT7/miR-21 binding and exosomal translocation in CECs. Hence, this study reveals evidence for crosstalk between SP-NK1R and ADO-ADORA2B signaling pathways in CECs, which regulates TUT7/miR-21 interaction followed by polyuridylation of miR-21 and subsequent recruitment of the TUT7/miR-21 complex to secreted exosomes. In conclusion, our study unravels an important cellular relationship between the SP-NK1R and ADO-ADORA2B in the colonic epithelial cells and a role of their crosstalk in the regulation of post-transcriptional 3’-end uridylation of miR-21 and selective enrichment of miR-21 into exosomes released by colonic epithelial cells.

## Materials and methods

All authors had access to the study data and had reviewed and approved the final manuscript.

### Antibodies and reagents

Reagents used are as follows: protease inhibitors cocktail (Cell Signaling Technology, MA, USA), Penicillin-Streptomycin (Cat.# P4333; Sigma-Aldrich), exosome-depleted fetal bovine serum (Life Technologies, Carlsbad, CA), fetal bovine serum (Cat.# F2442; Sigma-Aldrich), M3D media (Incell, San Antonio, TX), advanced DMEM/F-12 (Cat.# 12,634–010; Invitrogen), DMEM (Cat.# D5796; Sigma-Aldrich), Y-27632 (Cat.# 1254; R&D Systems), SB 431542 (Cat.# 1614; R&D Systems), PSB-1115 (Cat.# 2009; Tocris Bioscience), siTUT7 and siControl (Ambion), Lipofectamine RNAiMAX (ThermoFisher), Matrigel (Cat.# 356234; BD Biosciences), anti-actin (1:1,000, A2066; Sigma Aldrich), anti-GAPDH (1:1,000, 5174S; Cell signaling Technology), anti-CD9 (sc-13118; Santa Cruz Biotechnology, Dallas, TX), anti-A33 (sc-398702; Santa Cruz Biotechnology), sCREB (1:1,000, 4820S; Cell signaling Technology), anti-p-CREB (1:1,000, 9198S; Cell signaling Technology), anti-pPKA (1:1,000, 5661S; Cell signaling Technology), anti-TUT7/anti-ZCCHC6 (1:500, 25196-1-AP; Proteintech).

### Generation of intestinal epithelial-specific NK-1R knockout mice

Mice bearing a LoxP-flanked NK-1R allele (NK-1R ^flox/+^ mice) were generated by inGenious Targeting Laboratory (Stony Brook, NY, USA). Briefly, a targeting vector was generated by inserting LoxP sites between flanking exon 2. Two independent NK-1R ^flox/+^ ES clones were identified, which were injected into C57BL/6 blastocysts to generate chimeric mice. The chimeric mice were bred with wild type C57BL/6 mice for germline transmission. Wild-type and NK1R^-/-^littermates were derived from subsequent heterozygous breeding in the Division of Laboratory Animal Medicine at UCLA. Heterozygous mice were then crossed with mice expressing flp-recombinase in the germline (Flipper mice, from Jackson lab, USA) to delete the FRT-flanked Neo cassette. Offspring of these mice were heterozygous for the desired NK-1R ^flox/+^ allele. Intestinal epithelial-specific NK-1R knockout mice were generated by crossing with Vil1-Cre (Jackson lab, B6.Cg-Tg(Vil1-cre)1000Gum/J (Stock no. 021504). 8-12 week-old male C57BL6 mice (n=8/group) were injected intraperitoneally and intracolonically with SP (72μg/mouse, 100μl; Bachem, King of Prussia, PA) or vehicle (0.1% trifluoroacetic acid), twice a day (8 hours apart on days 1 and 2). Male mice carrying wild type NK-1R (Vil1-cre;NK1R^+/+^) from the same litters were used for the control group. Mice were housed two to four per cage, maintained on a 12:12 hour light-dark cycle, and given access to food and water *ad libitum*. After all *in vivo* experiments, mice were euthanized on day 3 and the distal colons were removed for further study. In addition, colon tissues were collected for harvesting colonic epithelial cells and intestinal stem cells as previously described^48^. All mouse-based experiments were done in accordance with the UCLA Institutional Animal Care and Use Committee (IACUC).

### Cell lines, transfection and pharmacological treatment

Human colonic epithelial cells (NCM460 and NCM460-NK1R)^43^ were maintained in M3D media (Incell, San Antonio, TX) supplemented with 10% (vol/vol) exosome depleted FBS, 1% L-glutamine, 10 U/ml penicillin, and 100μg/ml streptomycin at 37°C, 5% CO2. Unless otherwise indicated, NCM460-NK1R cells were treated with SP (100nM, 6 hr, n=8/group). For TUT7 gene-silencing, NCM460-NK1R cells were transfected with siTUT7 (Ambion), using lipofectamine RNAiMAX (Invitrogen). Cells transfected with sicontrol (scrambled) served as controls (n=8/group). All transfections were performed 48 hr prior to SP stimulation. Pharmacological inhibition studies included treatment of NCM460-NK1R cells with A2B receptor antagonist, (PSB 1115: 1 mg/kg, Tocris Bioscience) (100nM, 1hr, n=8/group) or vehicle (sterile water).

### Culture of L-WRN Cells and production of L-WRN conditioned medium

L-WRN cells (ATCC®; Cat.# CRL-3276™) were cultured in high glucose DMEM supplemented with penicillin (100 units/mL), streptomycin (0.1 mg/mL) (Sigma-Aldrich) and 10% fetal bovine serum (Sigma-Aldrich). The 50% L-WRN CM was prepared with a 50/50 mix of the CM and fresh primary culture media, which is Advanced DMEM/F-12 (Invitrogen) supplemented with 20% fetal bovine serum (Sigma-Aldrich), 2 mM l-glutamine, 100 units/mL penicillin and 0.1 mg/mL streptomycin and also with 10μM Y-27632 (ROCK inhibitor; R&D Systems) and 10μM SB 431542 (TGF-βRI inhibitor; R&D Systems).

### 3D spheroid cell culture

Primary colonic epithelial stem cells were isolated from mouse colon (wild type, Vil1-cre;NK-1R^+/+^and intestinal-epithelial-specific NK-1R-Knock Out, Vil1-cre;NK1R^fl/fl^) and grown and maintained as 3D spheroid cultures in Matrigel (BD Biosciences) as described by Miyoshi et al^48^. Cells were kept in 50% L-WRN CM. Media were changed every 2 days, and cells were passaged every 3 days (1:3 split).

### Exosome isolation

Exosomes were isolated from the conditioned media of NCM460-NK1R cells (cultured in exosome-depleted FBS) treated with SP (100nM, 6 hr) or vehicle (0.1% TFA) by differential ultracentrifugation using Sorvall Discovery 90SE Floor Ultra Speed Centrifuge, 50.4Ti rotor (Beckman Coulter). Isolation was performed by a modified version of previously described protocols^13,49^. Briefly, cell culture media was subjected to differential centrifugation of 1,000 x g (5 min, keeping supernatants), 27,000 x g (35 min, keeping supernatants) and 33,000 x g (4hrs, keeping pelleted exosomes). Exosome pellets were washed in phosphate-buffered saline (PBS) to eliminate contaminating proteins and centrifuged again at 33,000 x g (4hrs to ON, keeping pelleted exosomes). The PBS was removed, and the exosomes were immediately re-suspended in 50-100 μl of Buffer A (150mM NaCl, 10mM HEPES, pH 7.4, 1mM EGTA, 0.1mM MgCl2). Purified exosomes were then stored at −80 °C until use. All centrifugation steps were performed at 4°C. Protein quantification of the collected exosomes was performed using BCA method (Pierce™ BCA Protein Assay kit, ThermoFisher Scientific) and equal amount of exosomes were collected for downstream applications. Equal amount of exosomes were denatured in Laemmli lysis buffer and subjected to western blot analysis using antibody against the colonic epithelial exosome-specific protein, A33 as well as CD9 (well-known exosome surface marker)^2,50^.

### Mass Spectrometry

Exosomes isolated by ultracentrifugation were lysed in equal volume of radio-immunoprecipitation assay (RIPA) buffer (Cell Signaling Technology) containing protease inhibitor cocktail (Cell Signaling Technology) and quantified using bicinchoninic acid assay (BCA assay, Thermo Scientific). The normalized lysates were then subjected to mass spectrometric (MS) analysis performed on Q Exactive™ Plus Hybrid Quadrupole-Orbitrap™ Mass Spectrometer. 1.0 ul sample was injected to an ultimate 3000 nanoLC, which was equipped with a 75μm x 2cm trap column packed with C183μm bulk resins (Acclaim PepMap100, ThermoScientific) and a 75μm x 15cm analytical column with C182μm resins (Acclaim PepMap100, ThermoScientific). The nanoLC gradient was 3-35% solvent B (A=H2O with 0.1% formic acid: B= acetonitrile with 0.1% formic acid) over 100min and from 35% to 85% solvent B in 5min at flow rate 300nL/min. The nanoLC was coupled with a Q Exactive™ Plus Hybrid Quadrupole-Orbitrap™ Mass Spectrometer (ThermoFisher Scientific, San Jose, CA).The ESI voltage was set at 1.9kV, and the capillary temperature was set at 275oC. Full spectra (m/z350-2000) were acquired in profile mode with resolution 70,000atm/z200 with an automated gain control (AGC) target of 3×106. The most abundance 15 ions were subjected to fragmentation by higher-energy collisional dissociation (HCD) with normalized collisional energy of 25. MS/MS spectra were acquired in centroid mode with resolution 17,500 at m/z200. The AGC target for fragmentations are set at 2×104 with maximum injection time of 50ms. Charge states 1,7,8, and unassigned were excluded from tandem MS experiments. Dynamic exclusion was set at 45.0s.

The MS data were analyzed using MaxQuant software version 1.3.0.5. MS data were used to query the Uniprot_HomoSapiens_154578_20160822.fasta. Raw data was searched again in uniprot human database by Proteome Discovered version 1.4. Following parameters were set: precursor mass tolerance ± 10ppm, fragment mass tolerance ± 0.02 Th for HCD, up to two miscleavges by trypsin, methionine oxidation as variable modification. False discovery rate was at 1.0% and minimum of 1 peptide was required for protein identification.

### RNA isolation and Real-time RT-PCR analysis

Total cellular as well as exosomal RNA were isolated using standard TRIzol reagent protocol (Ambion). Equal amount of total RNA from all cellular and exosome samples (200ng) were used to generate cDNA library using miRCURY LNA Universal RT microRNA PCR cDNA kit (Exiqon). miRNeasy kit was used according to each manufacturer’s total RNA isolation procedure. Quantitative RT-PCR (qRT-PCR) for miRNAs was performed using miRNA–specific primers (Exiqon) and miRCURY LNA Universal RT microRNA PCR SYBR Green master mix (Exiqon). The small non-coding RNA (snRNA) U6 (U6 snRNA-001973) and beta-actin were used as endogenous control. All quantitative reverse transcription polymerase chain reaction (RT-PCR) reactions were carried out in triplicates. The qRT-PCR for mRNAs of interest was performed using gene-specific primers (Applied Biosystems), according to the manufacturer’s instructions. The gene expression was calculated using the 2^-ΔΔCT^ value and the specificity of each reaction was confirmed by melting curve analysis. The RNA concentration was assessed using a NanoDrop 2000 spectrophotometer (Thermo Scientific, Waltham, MA, USA).

### sncRNA isolation

The small noncoding RNAs (sncRNAs) were separated from the NCM460-NK1R cells and their secreted exosomes by employing NucleoSpin® miRNA (MACHEREY NAGEL) Isolation Kit as per the manufacturer’s instructions. These sncRNAs were further processed for cDNA synthesis employing miScript Reverse Transcriptase Kit as per supplier’s protocol^16^. Subsequently, the intrinsic expression of miR-21 and miR-1307-3p within these cells and the amount of these miRNAs recruited to their secreted exosomes was quantified by qRT-PCR with gene-specific primers. The snRNA U6 was used as an invariant control for normalizing the expression of miR-21 and miR-1307-3p. The 2^-ΔΔCT^ was used to calculate the relative levels of both the miRNAs.

### Protein isolation and immunoblotting

Total cellular and exosome proteins were isolated with RIPA buffer (Cell Signaling Technology) containing protease inhibitors (Cell Signaling Technology). Protein quantification of the lysed cells and exosome samples was performed using the bicinchoninic acid assay (BCA assay, Thermo Scientific). Equal amount of lysates were resolved by sodium dodecyl sulfate polyacrylamide gel electrophoresis (SDS-PAGE) and transferred onto polyvinylidene fluoride membrane (Merck Millipore, Burlington, MA, USA) in 25 mmol/L Tris, 192 mmol/L glycine. Membranes were blocked (phosphate-buffered saline, 10% non-fat dry milk, 0.05% Tween-20) and probed with antibodies followed by corresponding horseradish peroxidase–labelled secondary antibodies (1:1000). Blots were developed with enhanced chemiluminescence reagent (SuperSignal™ West Pico PLUS Chemiluminescent Substrate, Thermofisher Scientific) and imaged by the ChemidocTM gel imaging system (Bio-Rad). Western blot bands were quantified by densitometry using ImageJ software (http://rsb.info.nih.gov/ij). Data are represented by cropped images from the original membranes. Glyceraldehyde 3-phosphate dehydrogenase (GAPDH) (Cell Signaling Technology) and anti-actin (Sigma Aldrich) were used as a loading control. Results for immunoblotting are representative of at least three independent experiments.

### Bioinformatic analysis

A potential interaction of TUT7 and miR-21 was assessed *in silico* by use of the RNA-protein interaction prediction tool (RPISeq), which is freely available online: http://pridb.gdcb.iastate.edu/RPISeq/references.php51. For the analysis, raw values predicted by the SVM classifier were used with a cut-off value of 0.5. JASPAR (http://jaspar.genereg.net/matrix-clusters/)52, TRANSFAC^53^, TFSEARCH^54^, databases of transcription factor binding models, were used to perform searches for putative transcription factor binding sites in DNA sequences.

### Quantification and statistical analysis

GraphPad Prism Analysis software (GraphPad Prism 9.0.2(161) Software, La Jolla, CA) was used to perform statistical analysis. Data are presented as the mean ± SD unless otherwise specified. All experiments were performed at least three independent times, unless otherwise noted. Student’s t-tests were used to evaluate significant differences between any two groups of data; one-way analysis of variance was used if there were more than two groups. Statistical significance was marked with ns, nonsignificant, * for *p* < 0.05, ** for *p* < 0.01, *** for *p* < 0.001 and **** for *p* < 0.0001. *p* < 0.05 was considered statistically significant.

### Physiological significance

SP/NK1R signaling activation promotes a diverse array of molecular signaling leading to cell proliferation, pro-inflammatory response, and exosome production in CECs in vitro and in mouse colitis models as well as in patients with IBD. Previously, our group has shown that SP/NK1R signaling activation promotes miR-21 production and preferentially exports it to the exosomes released by CECs. In the present study we unravelled that miR-21 exosomal export is regulated by the SP-induced activation of colonic epithelium-specific Adenosine A_2B_ receptor signaling cascade which promotes TUT7/miR-21 binding and subsequent TUT7-mediated polyuridylation of miR-21. This facilitates the selective sorting of miR-21 into the secreted exosomes from CECs. Moreover, Adenosine A_2B_ receptor signaling cascade is identified to have a key regulatory role in the progression of IBD, especially associated with ulcerative colitis (UC), and hence can be a pharmacological target for the therapeutic benefits in UC.

## Abbreviations

AC: adenylyl cyclase
ADO: adenosine
ADORA2B: adenosine A_2B_ receptor
ARC: animal research committee
AKT: protein kinase B (PKB)
cAMP: 3’,5’-cyclic adenosine monophosphate
CECs: colonic epithelial cells
(CEEs): colonic epithelial exosomes
CM: conditioned medium
CD: crohn’s Disease
CREB: cAMP response element-binding protein
CXCL1: C-X-C Motif Chemokine Ligand 1
ENT2: equilibrative nucleoside transporter2
GPCRs: G protein-coupled receptors
IBD: inflammatory bowel disease
IECs: intestinal epithelial cells
IGFR: insulin-like growth factor receptor
IL8: interleukin 8
MAPK: mitogen-activated protein kinase
miRNA: microRNA
MVB: multivesicular body
ncRNAs: non-coding RNAs
NF-κB: nuclear factor kappa-light-chain-enhancer of activated B cells
NK-1R: neurokinin-1 receptor
PKA: protein kinase A
RBPs: RNA binding proteins
ROCK inhibitor: rho-associated protein kinase inhibitor
RT-PCR: real-time polymerase chain reaction
sncRNAs: small non-coding RNAs
SP: Substance P
TENTs: terminal nucleotidyl transferases
TFA: trifluoroacetic acid
TGF-βRI: transforming growth factor-β type I receptor
TUTs: terminal uridyl transferases
UC: Ulcerative colitis
ZCCHC6: zinc finger
CCHC: domain containing 6

## Acknowledgments

We would like to thank UCLA Proteomic Core for their help with proteomic analysis. LC-MS was supported by an NIH shared instrumentation grant 1S10OD016387-01.

